# Identification of Stress-induced miRNA of Microbial flora, networking, expression with chronic rhinosinusitis (CRS)

**DOI:** 10.1101/2024.11.11.622768

**Authors:** Bo Wei, Jiaoyu He, Jiaxin Li, Junming Xian, Feng Lius

## Abstract

All age groups are affected by one of the most prevalent chronic medical disorders globally: chronic rhinosinusitis (CRS). Its estimated incidence is 10.9% in Europe, 13% in China, and 12.3% in the United States. For many years, the 16S rRNA gene has been the backbone of sequence-based bacterial study. Still, it is only recently that the possibility of high-throughput sequencing the entire gene has been feasible. A unique gene regulation mechanism known as “RNA silencing” limits the transcript level by either starting specific RNA degradation or blocking translation. RNA interference (RNAi) is commonly induced by viruses or exogenously produced small interfering RNAs. Small RNAs like these have been utilized in biomedical research to specifically silence genes. The purpose of a short hairpin RNA is to create interference by inserting an artificial RNA molecule with a hairpin or loop-like structure into a specially designed siRNA. Every stage of life, including cell development, metabolism, the cell cycle, and signal transmission, depends on gene regulatory networks. Total fifteen mRNAs of 16SrRNA gene examined, siRNA regions is unique and isolated through 3’ to 5’ position nucleotide. Networking of both genes lies varieties of path in human being, so early to utilized siRNA technique to silence the effect of 16SrRNA gene in CRS illness. The gene encoding 16S rRNA has around ten target sites that siRNAs can attach to. The siRNAs are made to be complementary sequences to each of the target sites. Verify the 16S rRNA gene expression in CRS illness and other types across the body. Since a cystic fibrosis transmembrane conductance regulator (CFTR) mutation causes CRS illness, we isolated a mutant form of CFTR with an accession ID of 1XMJ and 16S rRNA protein quotes in 8SR6. Examine the two proteins interaction (1xmj, 8sr6) describe the inhibition of CRS illness through new way. Utilizing in-vitro exploratory approaches, both this computational study of siRNA and protein-protein interaction may be employed further to verify efficacy and appropriateness.

## Introduction

All age groups are affected by one of the most prevalent chronic medical disorders globally: chronic rhinosinusitis (CRS). Its estimated incidence is 10.9% in Europe, 13% in China, and 12.3% in the United States (Albu, 2020). CRS is medical condition having decreased the quality of life of patients. Through literature it coded that CRS produced great effect on social functioning in heart and chronic heart failure (Hirsch et al., 2017). Moreover, research indicates that CRS costs society money as well. The direct expenses of CRS are thought to be between $10 and $13 billion USD annually in the USA. Additionally, absenteeism, lost productivity, and missed workdays are the indirect costs of CRS, which are projected to be more than USD 20 billion annually in the USA (Rudmik, 2017; DeConde & Soler, 2016). The Journal of Clinical Medicine (JCM) is widely known for publishing high-quality fundamental research and clinical investigations. Relevant papers for a wide readership are routinely published in the journal as a consequence of a rigorous peer-review process and meticulous selection procedure. The Prevention, Diagnosis and Management of Chronic Rhinosinusitis special issue of JCM is an example of the effort to present high-quality issues pertinent to researchers as well as physicians (Hu et al., 2019). Allergic illnesses, particularly IgE-mediated inflammatory processes like allergic rhinitis, are often believed to be a comorbidity/associated factor or an instigating factor for the development of CRS clinical state. The reasoning is that inflammation of the mucosa caused by allergies might result in occlusion of the sinus ostial passage and consequent secondary infection. Even though prior research has linked allergies to CRS, nothing is known about this connection (Marcus et al., 2018). For many years, the 16S rRNA gene has been the backbone of sequence-based bacterial study. Still, it is only recently that the possibility of high-throughput sequencing the whole gene has been feasible. Here, we critically reevaluate the 16S gene’s capacity to give taxonomic resolution at the species and strain levels using in silico and sequence-based methods (Johnson et al., 2019). PCR-amplified 16S sequences have generally been grouped based on similarity to create operational taxonomic units (OTUs) since the advent of high-throughput sequencing. Representative OTU sequences are then compared with reference databases to estimate probable taxonomy. While 16S is strong and handy, its use has led to the historical presumption that sequences with more than 95% similarity belong to the same genus and sequences with more than 97% identity belong to the same species.(Winand et al., 2019; Boudewijns et al., 2006; Petti et al., 2005).

**Figure 1.**
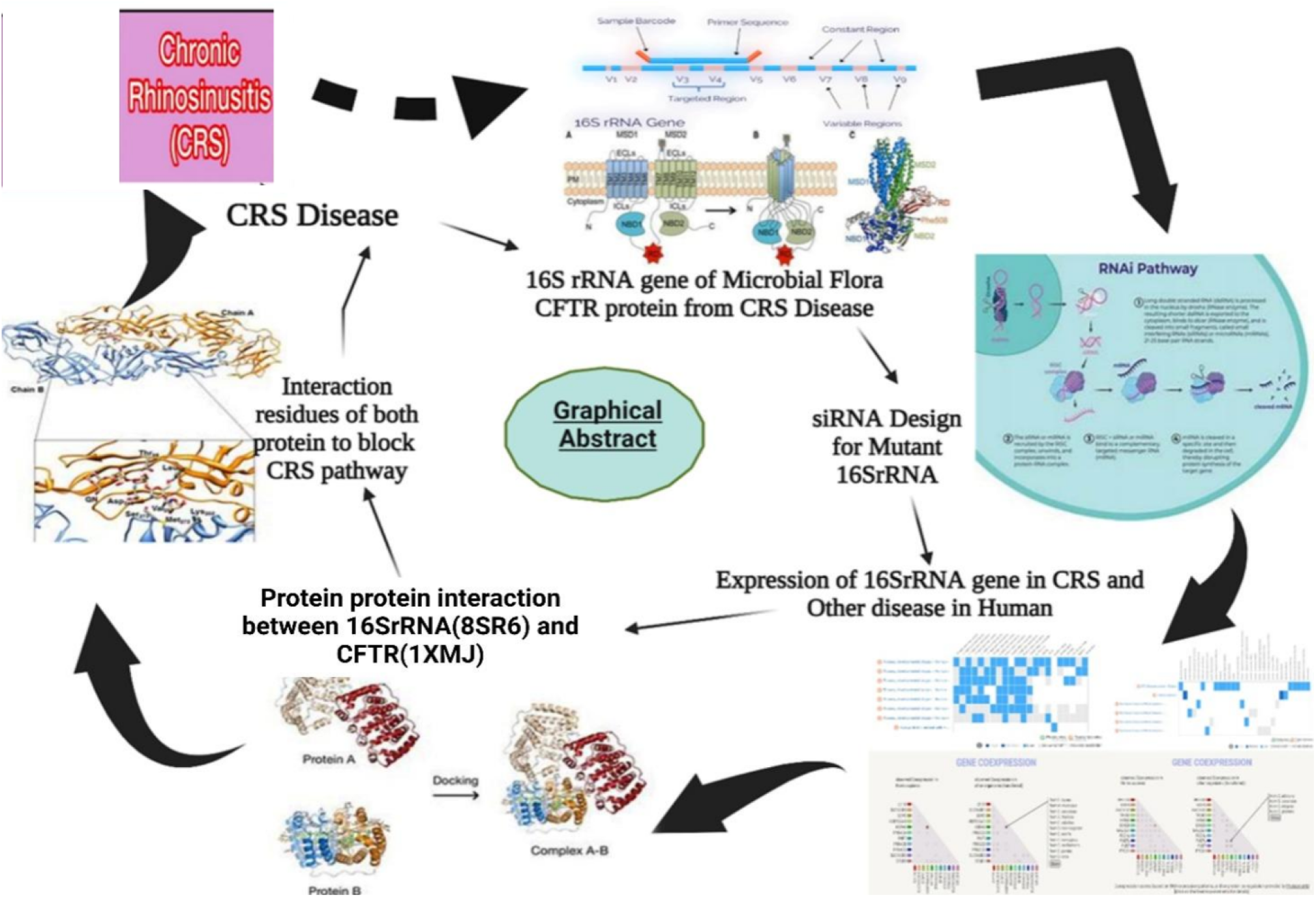
Illustrate the graphical abstract to follow methodology to Identification of Stress-induced miRNA of Microbial flora, networking, expression with Chronic Sinusitis

The field of microbiomics in chronic conditions, such as chronic rhinosinusitis (CRS), has advanced quickly in the last several years. Conceptually, high-impact microbiome research from other organ systems—like the gut—has been extended to the respiratory domain. Although 16S sequencing is a brand-new technique for studying microbes in the lab, recent research has only strengthened our present understanding of the bacteriology of CRS and added very little in the way of new clinical information. Constructing strong clinical trials targeted at reversing the damaged CRS in addition to combining 16S investigations with more intricate NGS techniques. Methods for sequencing the whole and partial 16S rRNA gene have become effective tools for detecting bacteria with abnormal phenotypes (Anderson et al., 2016; Park et al., 2021; Jervis Bardy & Psaltis, 2016).

Every stage of life, including cell development, metabolism, the cell cycle, and signal transmission, depends on gene regulatory networks. We may learn more about the mechanisms behind disorders that result from dysregulation of these cellular activities by comprehending the dynamics of these networks. Biotechnological initiatives will advance more quickly if regulatory network behavior can be accurately predicted, as these predictions can be made more quickly and affordably than lab tests. Computational techniques have previously shown to be an effective research tool for assisting in the creation of network models as well as for analyzing their functionality (Karlebach & Shamir, 2008). Strategies linked to RNA interference have gained popularity as research approaches in a number of domains. A crucial stage in the silencing process is precisely designing the sequence of these tiny molecules. Many scholars have attempted to design certain methods to improve the likelihood of small interfering RNAs (siRNAs) becoming successful. Online designing software has been developing siRNA designs based on the most recognized algorithms in the last few decades with the goal of improving their quality (Fakhr et al., 2016).

Each strand of siRNA has 2-to 3-nucleotide overhangs as well as 5-phosphate and 3-hydroxyl endings. Remarkably, exogenously administered siRNAs that were generated in vitro may cause specific RNA degradation when introduced to distinct cells(Elbashir, Lendeckel, et al., 2001). In several vertebrates and invertebrates, these siRNAs have also been observed to specifically impede gene expression(Zhang et al., 2024). The direct biochemical validation that siRNAs may function as guide RNAs for associated mRNA degradation has been provided recently by Schwarz et al (Schwarz et al., 2002). Similarly, small hairpin RNA (shRNA) is an artificially manufactured RNA molecule with a constricted hairpin turn, utilized to inhibit the target gene expression by RNA interference (Brummelkamp et al., 2002; Paddison et al., 2002). SiRNA expression in cells can be achieved by plasmid delivery or via viral or bacterial vectors. Transfection of plasmids to cells using transfection to gain siRNA expression may be accomplished with economically accessible chemicals in vitro (Xiang et al., 2006). Until recently, nothing has been done to create siRNA and shRNA for the Microbiome 16S rRNA gene expression function in CRS. This study effort attempts to computationally develop siRNA of gene 16S rRNA function, expression, and analysis in the illness CRS.

## Material and Methods

### Literature review and Sequence Retrieval

The NCBI database, which can be accessed at https://www.ncbi.nlm.nih.gov, contains all of the information about the mutations of the Microbiome 16S rRNA gene, as well as the mutated mRNA sequences of the Microbiome 16S rRNA gene with accession ids FM208770.1, FM208766.1, FM20867.1, FM208768.1k, FM208769.1, FM208775.1, FM208776.1, FM208777.1, FM208778.1, FM208779.1, FM208780.1, FM208781.1, FM208782.1, FM208783.1, and FM208784.1. Concerned CRS disease protein was identified from the literature. The CFTR is the best receptor for CRS-related disorders. Almost all people with two CFTR mutations and cystic fibrosis (CF) will develop CRS. Both genes and proteins are reviewed from the literature.

### Identification of Start codon, target sites, UTR, and Expression of related gene and Proteins

The altered mRNA sequences were evaluated, and each mRNA sequence’s start codon (AUG) was discovered using the codon finder program(Hamady et al., 2009). The start codon is the first codon in the mRNA sequence, from which translation begins. For siRNA and shRNA binding to the altered 16S rRNA gene, the targeting sites sequence motifs on the gene are discovered using the siRNA Target Finder provided at https://www.genscript.com (Lück et al., 2019). 5 to 3 untranslated regions (UTRS) were found using RegRNA service(Huang et al., 2006). Tandem repeats were detected using the Tandem Repeats Finder site (Benson, 1999). The 3’ untranslated region (3’-UTR) is a section of mRNA that immediately follows the translation termination codon. An mRNA molecule is synthesized from the DNA sequence and then translated into a protein. The 3’-UTR has binding sites for both regulatory proteins and microRNAs. Expression of related gene of Microbiome from the BacwGSTdb 2.0 database (Ruan & Feng, 2016) and CRS disease CFTR protein from the gene cards (Stelzer et al., 2016).

### Homology, siRNA design, prediction of structures

To locate similar gene sequences, a homology search was performed on each mRNA sequence’s target site sequence using the NCBI Blast database. It is critical to avoid homologous target sites because if a similar sequence appears in any gene as a target site, the intended siRNA or shRNA may attach to the normal gene’s target site and inhibit its production. The siRNA were created for each target site sequence with the help of the OligoWalk service(Lu & Mathews, 2008). Each siRNA was analyzed, and siRNA Hairpins of varying lengths (3 to 9 nucleotides) were introduced into the two complimentary areas. The structural prediction of the 16S rRNA gene and CFTR protein using the Uniprot database.

### Mechanism Pathways and Protein protein Interaction

The KEGG database (http://www.kegg.jp/ or http://www.genome.jp/kegg/) is an encyclopedia of genes and genomes. The primary goal of the KEGG database project is to assign functional meanings to genes and genomes both at the molecular and higher levels. Molecular-level functions are stored in the KO (KEGG Orthology) database, where each KO is defined as a functional ortholog of genes and proteins (Kanehisa et al., 2017). One popular resource for protein–protein docking is the ClusPro server (https://cluspro.org). With just two files in Protein Data Bank (PDB) format needed for basic usage, the service offers a straightforward home page. The removal of unstructured protein regions, the application of attraction or repulsion, the accounting for pairwise distance constraints, the construction of homo-multimers, the consideration of small-angle X-ray scattering (SAXS) data, and the location of heparin-binding sites are just a few of the advanced options that ClusPro offers to alter the search. Examine the two proteins’ control, interaction, and accessibility (Kozakov et al., 2017).

### Analysis and Interpretation

We provide a graphical approach that uses 3D coordinates to automatically create several 2D diagrams of ligand-protein interactions. The diagrams show the hydrophobic interactions and hydrogen-bond interaction patterns between the ligand(s) and the protein’s main or side chains. The system may depict linked sets of ligand-protein interactions in the same direction(Laskowski & Swindells, 2011).

**Figure 2.**
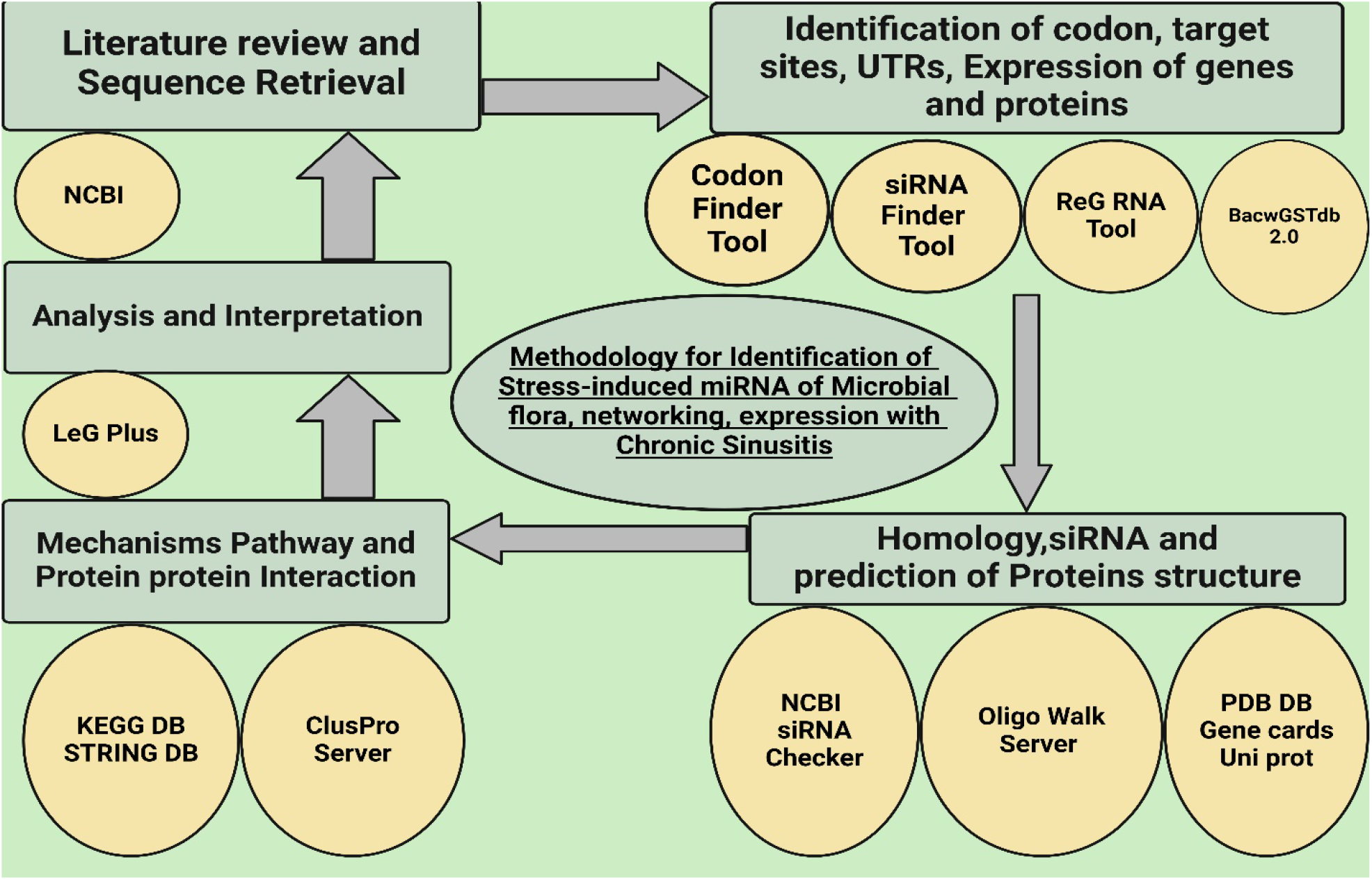
Schematic methodology for Stress induced siRNA, Identification of Microbiome, networking, expression in chronic rhinosinusitis (CRS)

## Results and Discussion

The selected mRNA sequences of 16S rRNA gene consist of fifteen variants. FM208770.1, FM208766.1, FM20867.1, FM208768.1, FM208769.1, FM208775.1, FM208776.1, FM208777.1, FM208778.1, FM208779.1, FM208780.1, FM208781.1, FM208782.1, FM208783.1 and FM208784.1. FM208770, FM 208766.1, FM208769.1, FM208775.1, FM208776.1, FM208777.1, FM208779.1, FM208780.1, FM208781.1, FM208783.1 variant 1 (16S rRNA partial) have no start translation frame AUG codons. FM20867.1 variant 2 (16S rRNA partial) has start translation frame AUG codons which help in translation. FM20868.1 variant 3 (16S rRNA partial) has start translation frame AUG codons which help in translation, direct exon extends. FM20878.1 variant 3 (16S rRNA partial) has start translation frame AUG codons which help in translation of protein. FM20882.1 variant 3 (16S rRNA partial) has start translation frame AUG codons which help in translation of protein. FM20882.1 variant 3 (16S rRNA partial) has start translation frame AUG codons which help in translation of protein. FM208770.1, FM208766.1, FM20867.1, FM208768.1, FM208769.1, FM208775.1, FM208776.1, FM208777.1, FM208778.1, FM208779.1, FM208780.1, FM208781.1, FM208782.1, FM208783.1 and FM208784.1 all mRNAs have not occupied any repetition in his sequence. It’s accomplished that all mRNA have cDNA regions available. Next we find siRNA target from all the mRNAs. Launch the codon every mRNA sequence started with AUG. After searching each mRNA sequence for target sites, the AA dinucleotide and the 19 3’ surrounding nucleotides were selected as important siRNA target sites. The belief of Elbashir et al. that siRNAs with 3’ overhanging UU dinucleotides are the best and successfully activate RNAi is the basis for this process for selecting siRNA target sites. This is also useful for translating and transcribed siRNAs using RNA pol III since RNA pol III terminates translation at 4-6 nucleotide poly(T) tracts, resulting in RNA particles with a short poly(U) tail(Elbashir, Martinez, et al., 2001). The same target sites were found in each mRNA transcript sequence; table 1 displays all of the target sites and the G+C content ratio. Ten siRNAs with a GC% ratio in the sense and antisense strand regions are present in all mRNAs. Every mRNA has a GC ratio of 50% or above, with beginning regions revealed as well.

**Table 1.**
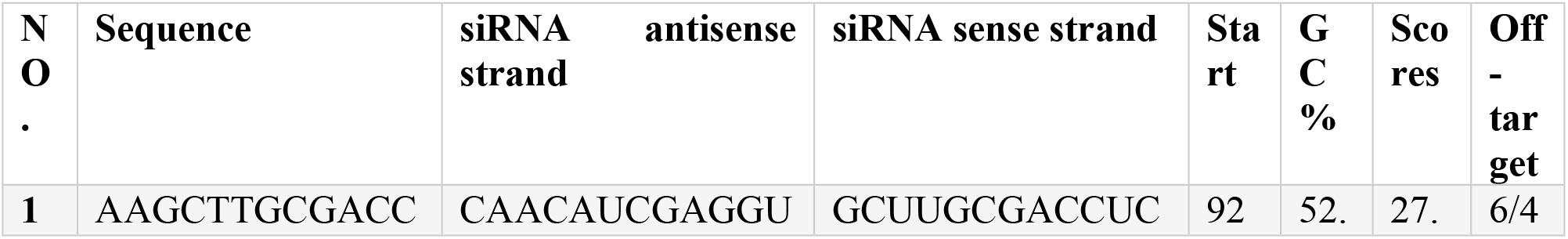
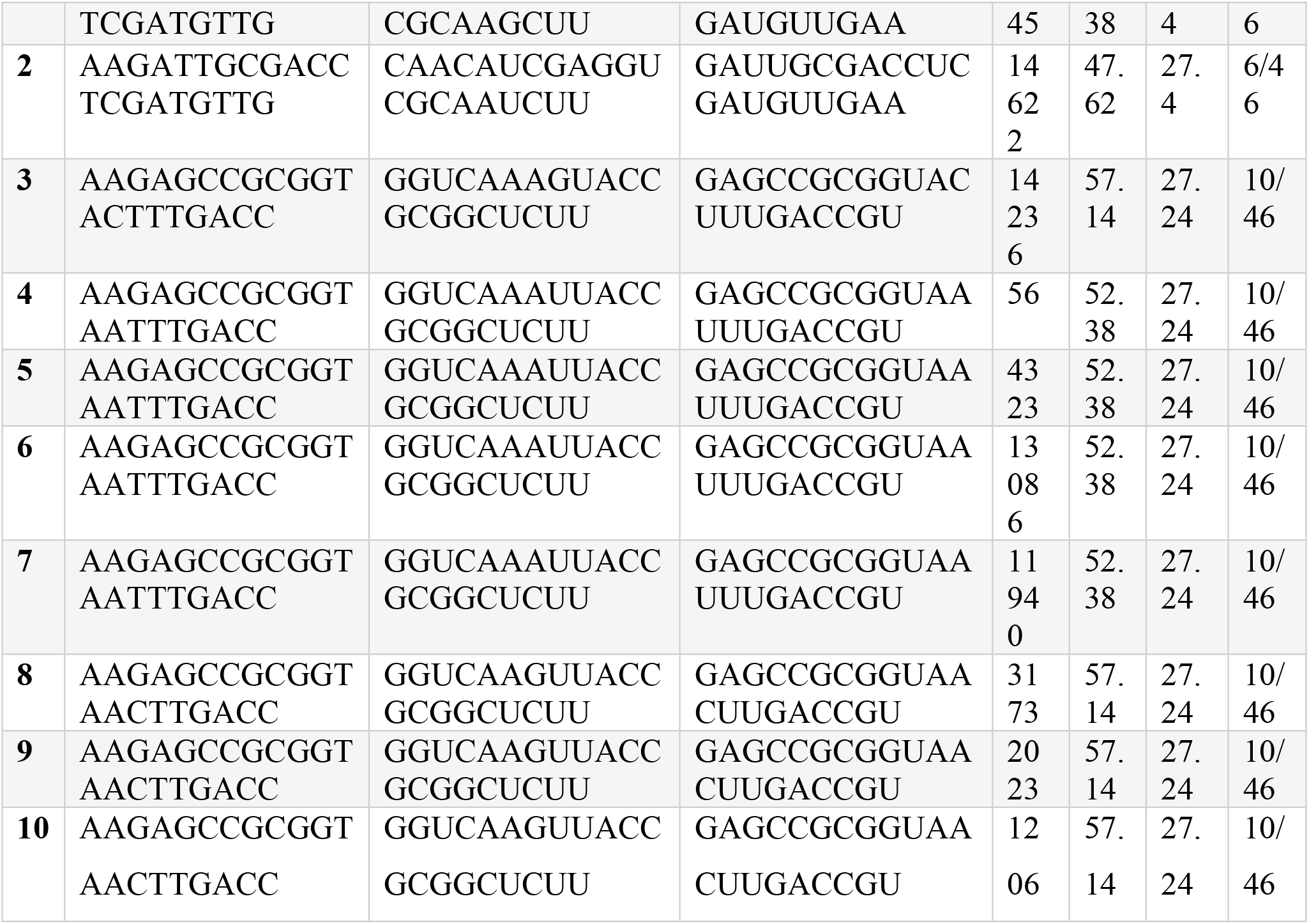
List of all mRNAs sequence siRNA targets having antisense and sense strand ratio with GC percentage.

SiRNAs with 30–50% G+C content are more energetic and more likely to operate as siRNAs than siRNAs with a higher G+C content percentage. 5’URT and 3’UTR should be avoided while designing siRNAs, despite the fact that UTRs that target siRNA have been shown to successfully induce gene silence. The section of the translation end codon that immediately follows is known as the 3’ untranslated region. A few mRNA sections, including as the 5’ UTR, 5’ cap, poly (A) tail, and 3’ UTR, are not translated into proteins. There are frequently regulatory areas in the 3’-UTR that affect the gene’s expression post-transcriptionally. The 3’UTR has a median length of 700 nucleotides. The 3’UTR contains regulatory regions that can affect the mRNA’s polyadenylation, translation efficiency, stability, and localization (Barrett et al., 2012;Pichon et al., 2012). The expression of the 16S rRNA gene in human illness and the incidence of tissue development are described in Figure 3. The most likely gene that humans are born with that contributes to the spread of various cancers, cell division, and destruction capacity. The translation of 16SrRNA gene into various components, its help to increase bacterial growth in homosapiens, especially concerned in pneumonia alvoli section 8SR6 protein involved. The structure of the CFTR protein, a key component of CRS illness, is derived from Uniprot RSCB PDB database(1XMJ). 1XMI have major component of CFTR protein complex, having mutational chains available in it. 1XMJ is same derivation year and having x-ray crystallography ratio is 2.3 contain single chain. Alpha helices and beta sheets in the structure include appealing amino acids with strong binding affinities. The area of the mRNA directly upstream of the initiation codon is known as the 5′ UTR. This area is involved in transcription control. Each mRNA transcript’s comparable untranslated sections and corresponding positions were found. Tandem repeat elimination is a crucial step in the discovery of siRNA target locations. Typically, a tandem repeat consists of two or more neighboring copies of the same nucleotide sequence. The target site will not bind to siRNA if it ends with a length of four or five repetitions. Repeats are thus often eliminated throughout the siRNA design process. The 16S rRNA gene’s mRNA straps are all unique and include no repeat, which calls for more investigation.

**Figure 3.**
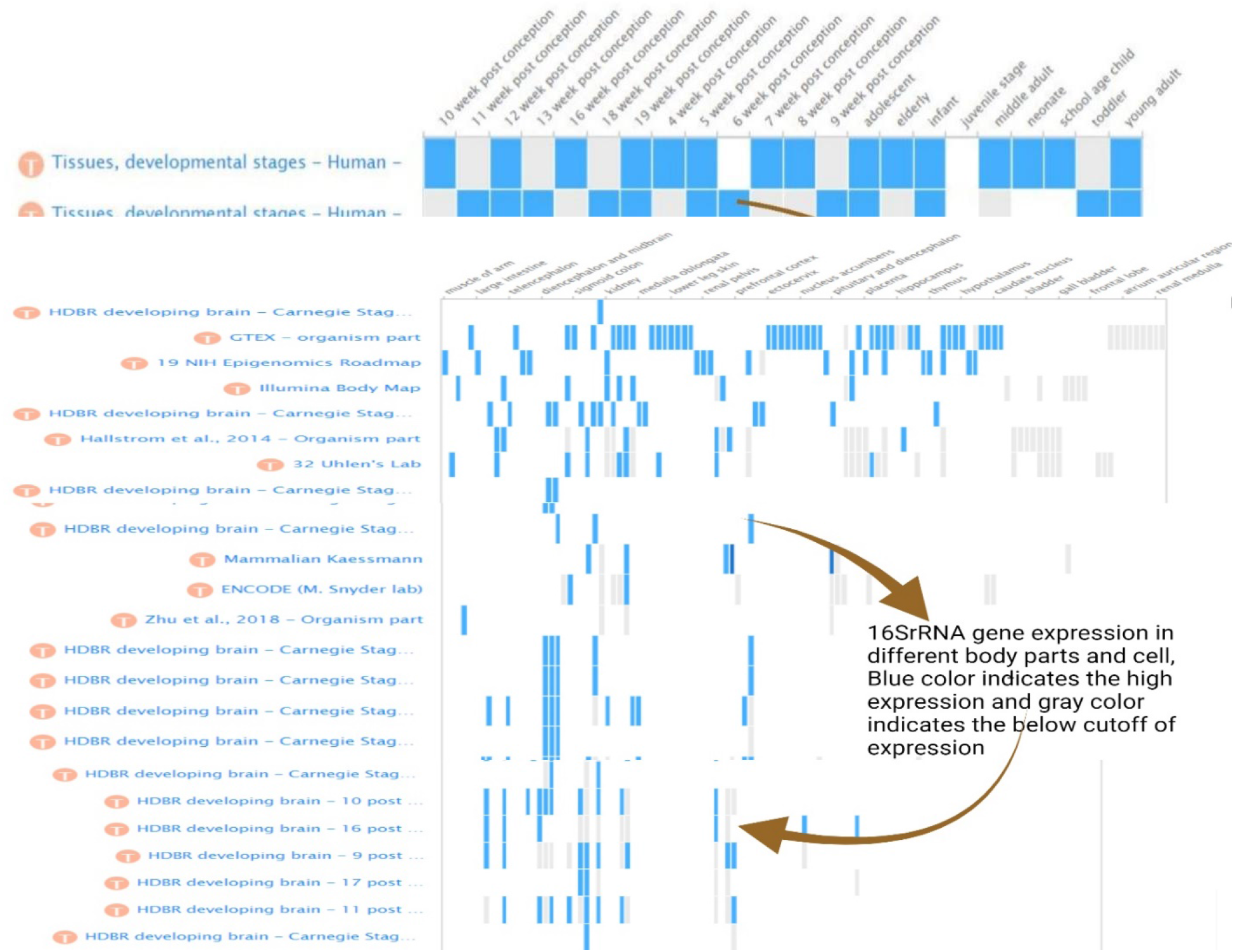
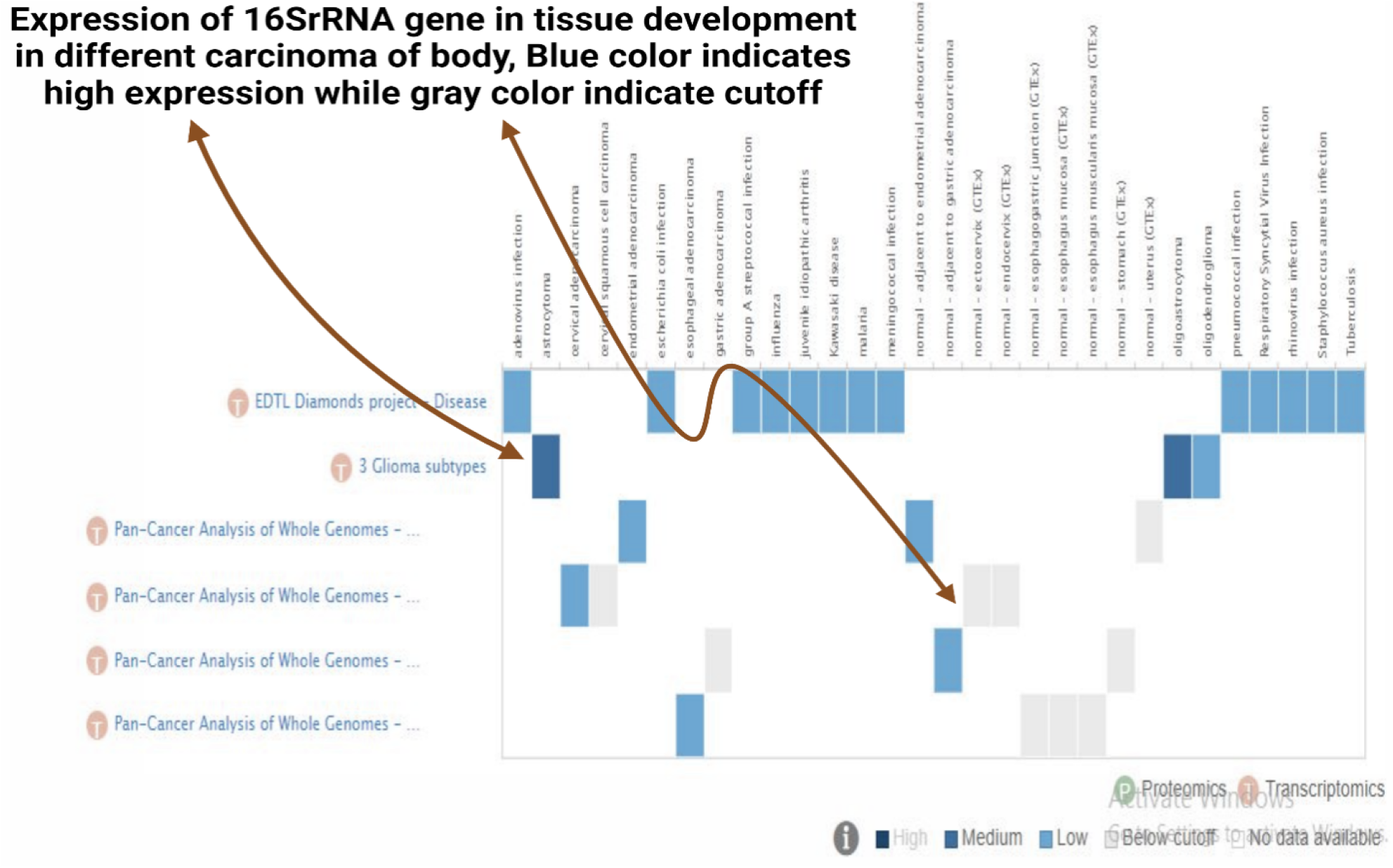
Illustrate the expression of 16SrRNA gene in different carcinomas, regions of the body and genome state of the body, Blue color indicates the rich expression while gray color indicated the cutoff expression of gene

Every siRNA is a well-known and distinct one, although all mRNAs lack a homology search in the NCBI. Then, we investigate the structural feature, CFTR, and 16S rRNA in the homology search; nevertheless, none of the genes with comparable sequences that are identified as anticipated target sites. Avoiding homologous target sites is essential because there is a possibility that the siRNA or shRNA that is being created will bind to the target site of the normal gene and inhibit its production if a similar sequence is found in any gene that serves as a target site. Different criteria for siRNA design were used by experts who initially reported using siRNA expression vectors to induce RNA interference. Two inverted repetitions separated by a brief buffer sequence constituted a significant portion of the designs, which concluded with a string of Ts acting as the translation end site (Ge et al., 2010). The length of the nucleotide arrangement being used as siRNA varies to varying degrees. A few studies have employed the 19-nucleotide sequence as the siRNA (Miyagishi & Taira, 2002). On the other hand, some research teams have employed siRNA that ranges in length from 21 to 25–29 nucleotides (Murali et al., 2015). It is discovered that all siRNAs of these various lengths may activate gene silencing (Kutter & Svoboda, 2008). 19 nucleotides in length were used to design the siRNA used in this study; 15 distinct siRNA were created for each target site. For further details, see the top-ranked siRNA reselection. In Table 2, the siRNA are displayed. Table 3’s stated probability values for effective siRNA are excellent. The server’s overall positive predictive value is 0.954, which indicates that 78.6% of the siRNAs it chooses will effectively silence targets. Testing against a database of siRNA studies carried out under various experimental settings yields the positive predictive value.

**Table 2.**
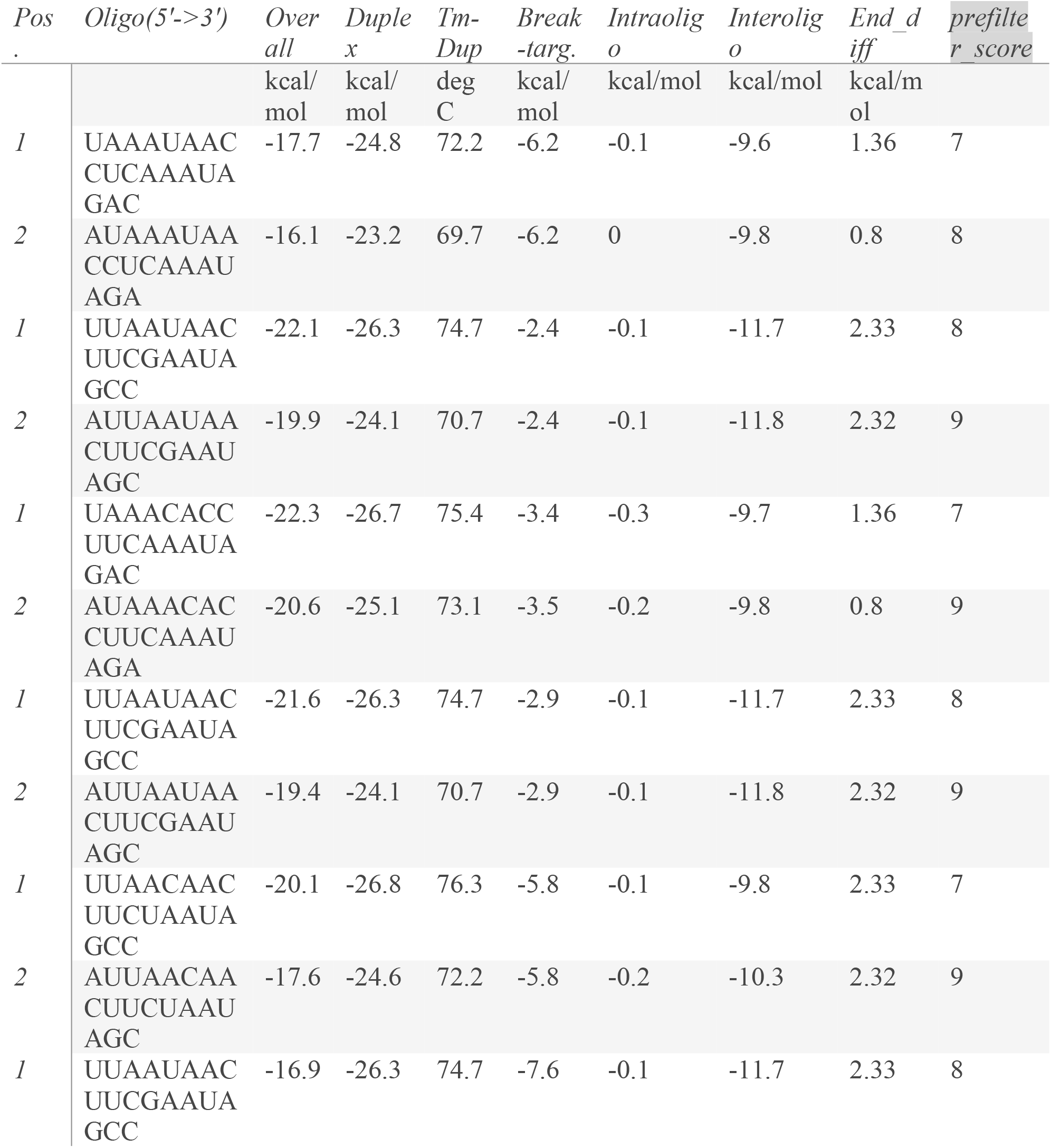
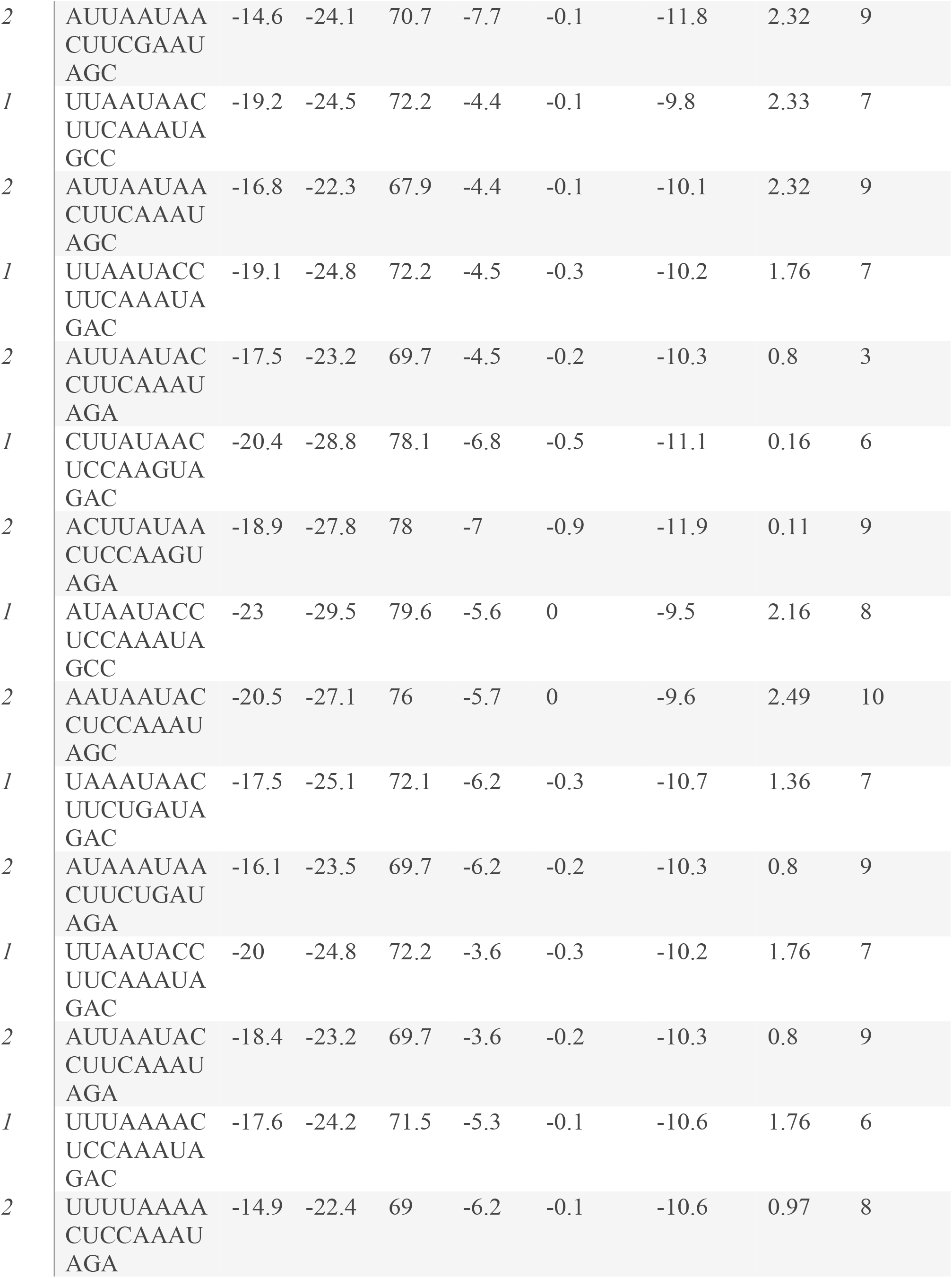
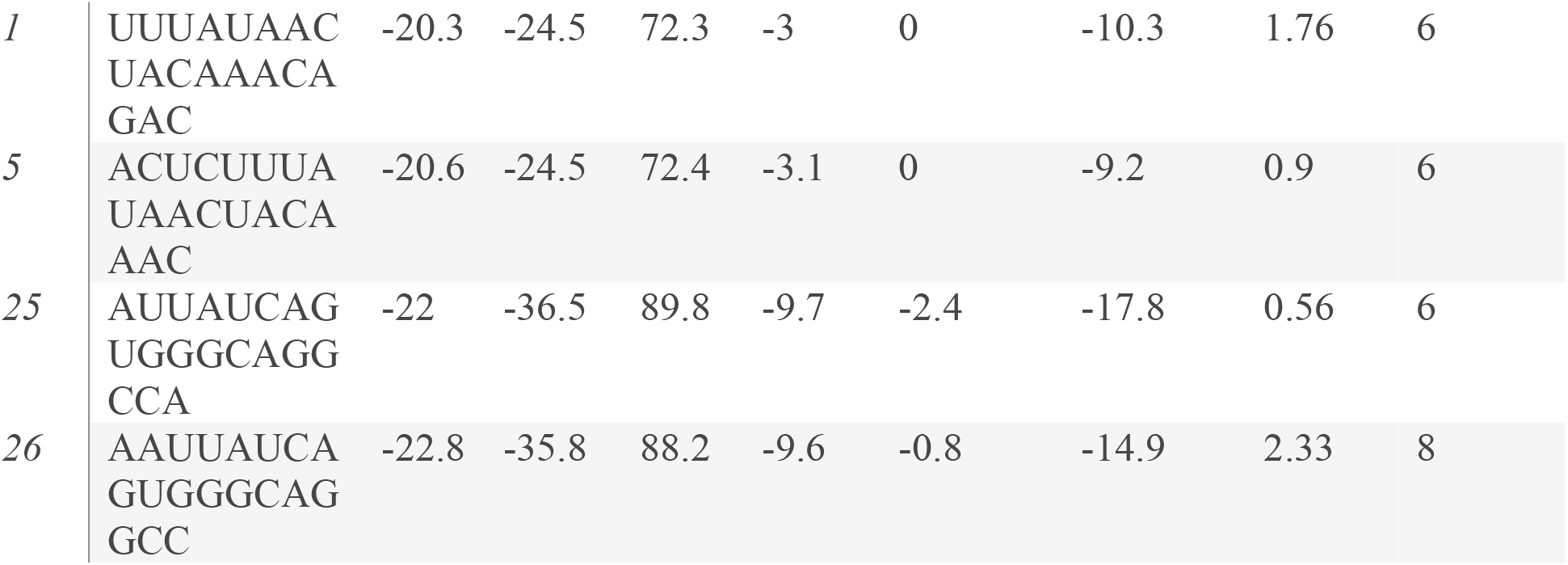
List of all siRNAs of predicted fifteen mRNAs. Having oligo position, Energy interoligo and intraoligo energies.

**Table 3.**
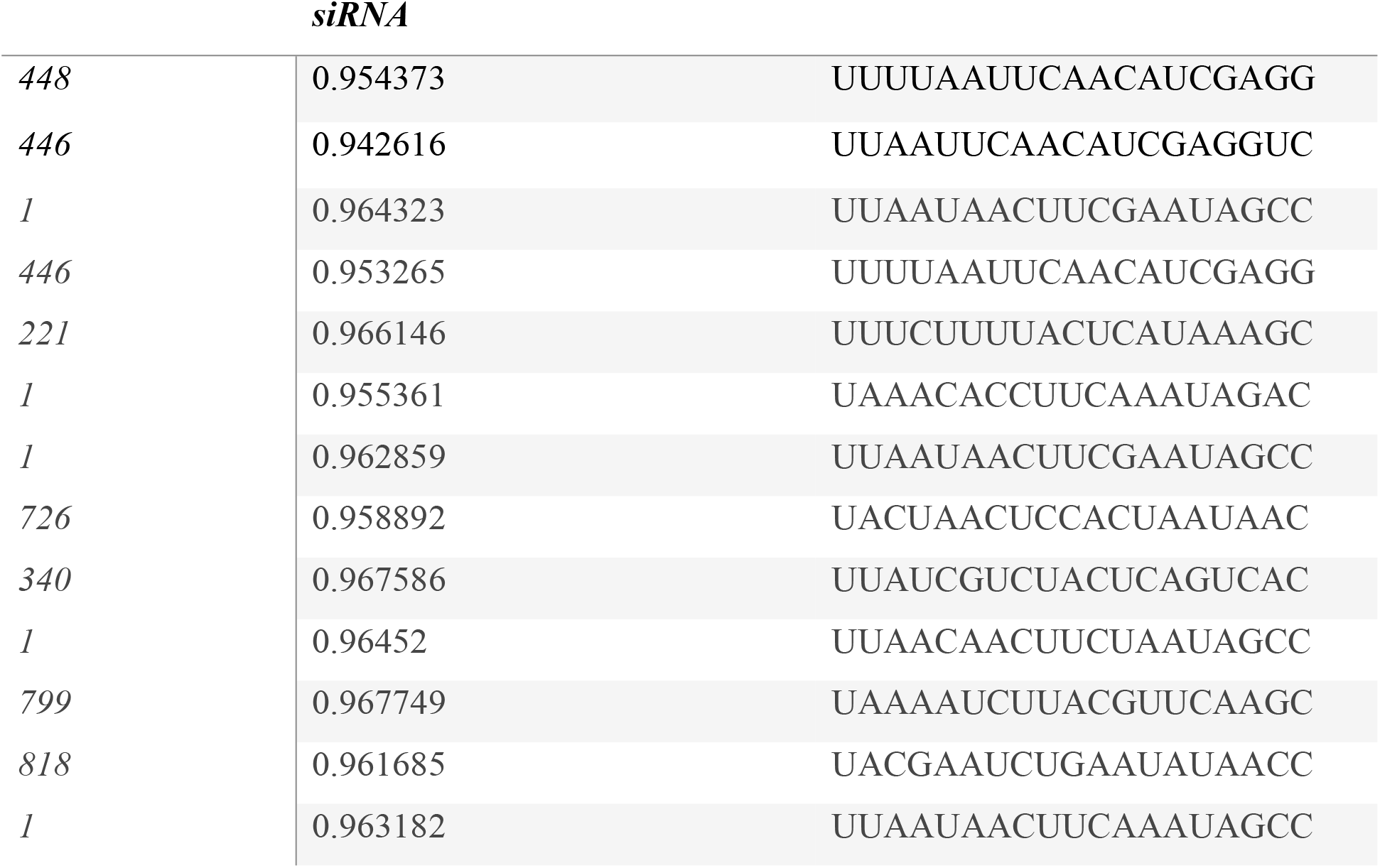
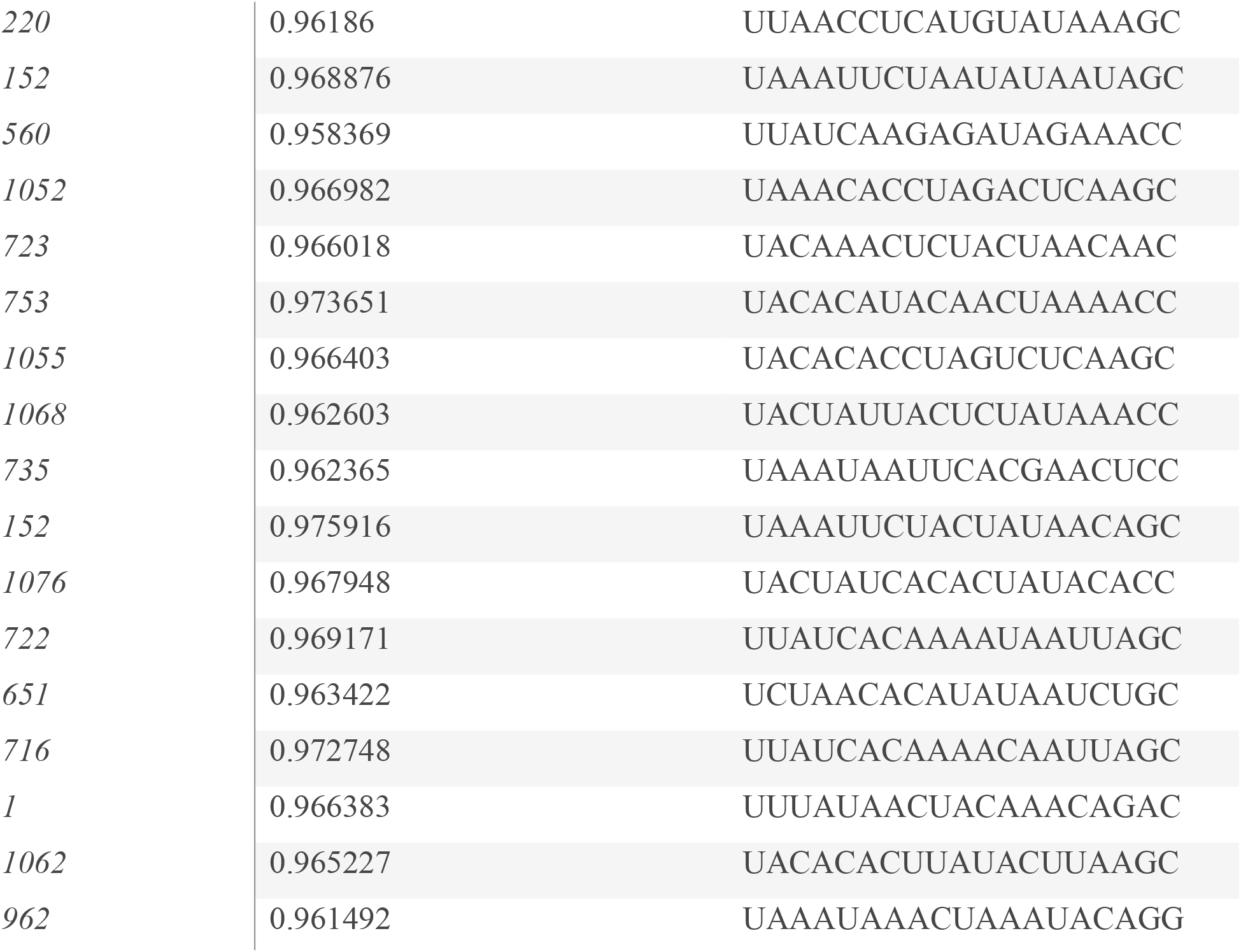
List of expected siRNAs having probability efficient score towards the target silencing.

Since all of the likelihood values in this instance are higher than 0.8, it is certain that the anticipated siRNAs will be more successful in silencing the mutant 16S rRNA gene. Several research teams have demonstrated successful hairpin siRNA gene silencing using a loop of three to twenty-three nucleotides (Penalva & Sánchez, 2003). According to our findings, all siRNAs have a probability larger than 0.8, indicating that they are a superior option for silencing a gene’s impact.

Typically, a hairpin siRNA is created with the target’s sense strand first, then its antisense strand in a certain order, separated by a brief spacer. For every target site sequence, a hairpin siRNA was created. If the planned hairpin construct does not start with a purine at siRNA exert 3, an additional ‘G’ is inserted to the construct’s beginning. It is preferred by RNA Poly III to use a purine to start transcription. Typically, siRNA transports the siRNA duplex directly to the cytosol, where it may bind DNA. It is composed of two complementary 19–22 bp RNA sequences joined by a brief loop of 4–11 nucleotides, akin to the hairpin present in actual miRNA. The protein 1XMI, which have seven chains and a structure between 2 and 2.3 xray crystallography score, is encoded to the CFTR protein complex that directly associate in CRS illness. Crystal structure of NBDI domain associated single chain A 1XMJ (Crystal structure of human F508A NBD1 domain with ATP). Both 1XMJ and 1XMI discovered in same year in 2005 for the component factor for CFTR in homosapiens. 1XMJ contain normal one chain taking for the further analysis, having structural conform changes Alpha helix, Beta sheets, coil and loop region avsilsble in it. The protein 8SR6 (Crystal structure of legAS4 from Legionella pneumophila subsp. pneumophila with histone H3 (3-17)peptide), which aids in translation, is directly responsive to 16SrRNA gene coding. The 8SR6 protein mutation directly affects the bacterial infection and help in spreading to all the body. The protein 8SR6 directly connect to Eukaryotic huntingtin interacting protein B in homosapiens which predicted in species Legionella pneumophila subsp. pneumophila. The structural features of the 1XMJ and 8SR6 proteins are explained in Fig 4.

**Figure 4.**
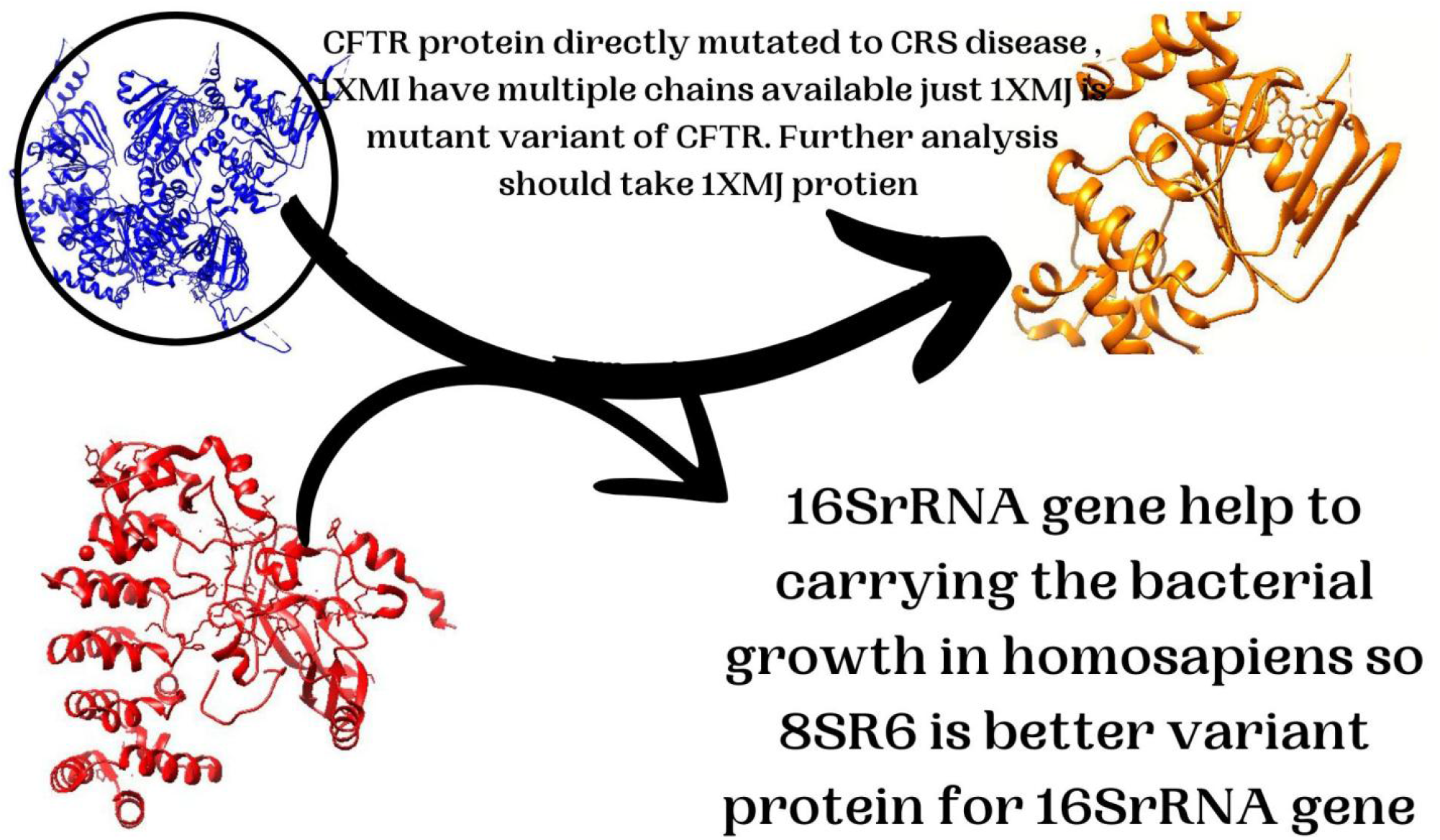
Illustrate the features and appearance of protein IXMJ having single chain available that retrieved from the IXMI multiple mutant protein of CFTR component hiving directly proportion to CRS illness, 16SrRNA gene translate into 8SR6 protein component

Our understanding of the molecular causes of human disease is nevertheless hampered by the persistent incompleteness of the data, despite extraordinary experimental efforts to map out the human interactome. An encouraging substitute is provided by computational techniques, which aid in the discovery of physiologically meaningful but as-yet-unmapped protein-protein interactions (PPIs). Although biological or network-based similarity is the basis for link prediction approaches to connect proteins, comparable proteins do not always interact and interacting proteins are not always similar. Here, we present structural and evolutionary evidence supporting the idea that proteins interact when one of them is similar to the partner of the other rather than when they are similar to one other. Ozger identified the SARS-Covid-2 network route, its interactions with other proteins, and its effects on other proteins(Kovács et al., 2019; Wang et al., 2023;Ozger, 2023). Our study looks into the various biological functions and pathways that the 16S rRNA (8SR6) protein participates in, such as mRNA pseudouridine synthesis, mitochondrial ribosome assembly, enzyme-directed rRNA pseudouridine synthesis, assembly of the large ribosomal subunit in the mitochondria, and positive regulation of mitochondrial translation. It culminated in rRNA binding, rRNA methyltransferase activity, pseudouridine synthase activity, and rRNA (guanosine-2-O-)-methyltransferase activity on a molecular level. It coordinated several pathways inside the cell, including the mitochondrial nucleoid, ribosome, matrix, granule of ribonucleoprotein, and mitochondrion.

The biological functions and pathways that the CFTR (1XMJ) protein is involved in vary. These include chaperone-mediated autophagy, protein kinase A signaling, high-density lipoprotein particle assembly, negative regulation of the smoothened signaling pathway involved in dorsal/ventral neural tube patterning, and regulation of anion channel activity. AMP-activated protein kinase activity, Type 2 metabotropic glutamate receptor binding, Type 3 metabotropic glutamate receptor binding, cAMP-dependent protein kinase activity, and TPR domain binding were the molecular outcomes. It synchronized many pathways with cAMP-dependent protein kinase complex, ciliary base, plasma membrane, and membrane in a biological manner. The co expression and network module of the CFTR protein (1XMJ) and 16S rRNA (8SR6) are described in Figure 5.

**Figure 5.**
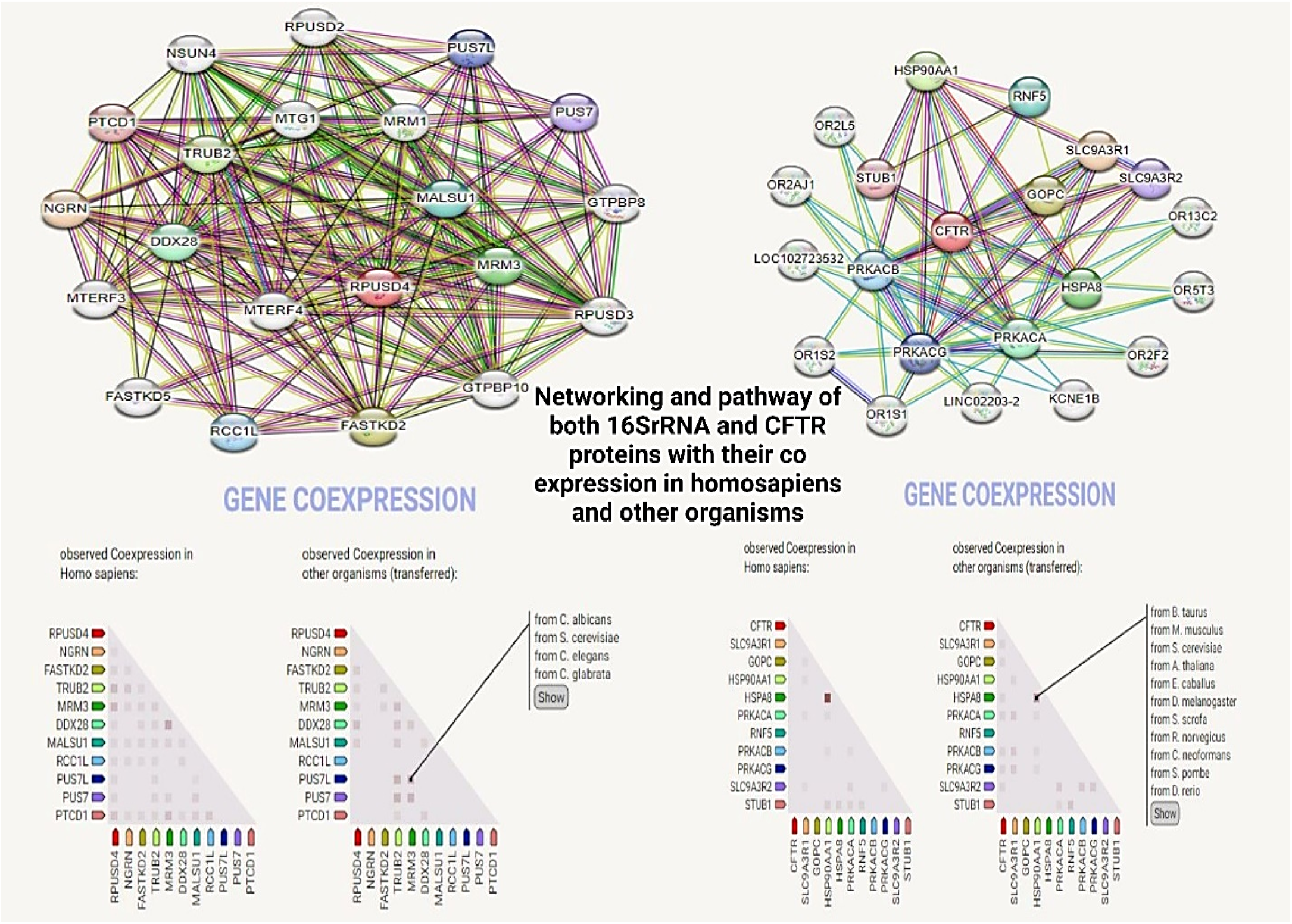
Explain the networking and pathway module of both 16SrRNA and CFTR proteins. Both proteins participate in various pathways and their co-expression lies in homosapiens and other organisms with great interest score

The scientific community became interested in studying the properties of proteins because of their biological relevance. The research provided insight into how proteins interact and serve many purposes in a live organism (Sunny & Jayaraj, 2022). Protein–protein interactions (PPIs) have a variety of vital functions in cells, including those of protein–protein inhibitors, antibody–antigen complexes, and super complexes. It is amazing how structural analysis techniques, such as cryo-EM, have advanced recently for determining protein complex structures (Tsuchiya et al., 2022). Protein–protein interactions (PPIs) have a variety of vital functions in cells, including those of protein–protein inhibitors, antibody–antigen complexes, and super complexes. Remarkable strides have been made recently in the identification of protein complex structures by structural analysis techniques, such as cryo-EM. When compared to C6 rat glioblastoma cells, naringin exhibits a greater cytotoxic potency against U-87MG human glioblastoma cells, suggesting that it may be used as a therapeutic treatment for glioblastoma (Uchikoga & Hirokawa, 2010). Our research concluded that accession key 1XMJ protein for CFTR and 8SR6 protein for 16SrRNA display docking interaction. The 8SR6 protein treat as a receptor having active residue, which bind to other protein with high affinity affinity; THR207;THR208;ASP115 and ARG113. Similarly 1XMJ protein treat as ligand in this complex shown their interaction residues; ASP443 and GLU632. Both proteins (ligand +Receptor Complex) interact each other with lowest energy -844.0 and -895.9 and bond lies at center of residues having distance noticed. This protein protein interaction complex cluster the residues of both proteins having distance 2.60, 3.00,2.89 and 2.46. Figure 6 illustrate the docking interaction, hydrophobic and distance between the bonds of interacting residues.

**Figure 6.**
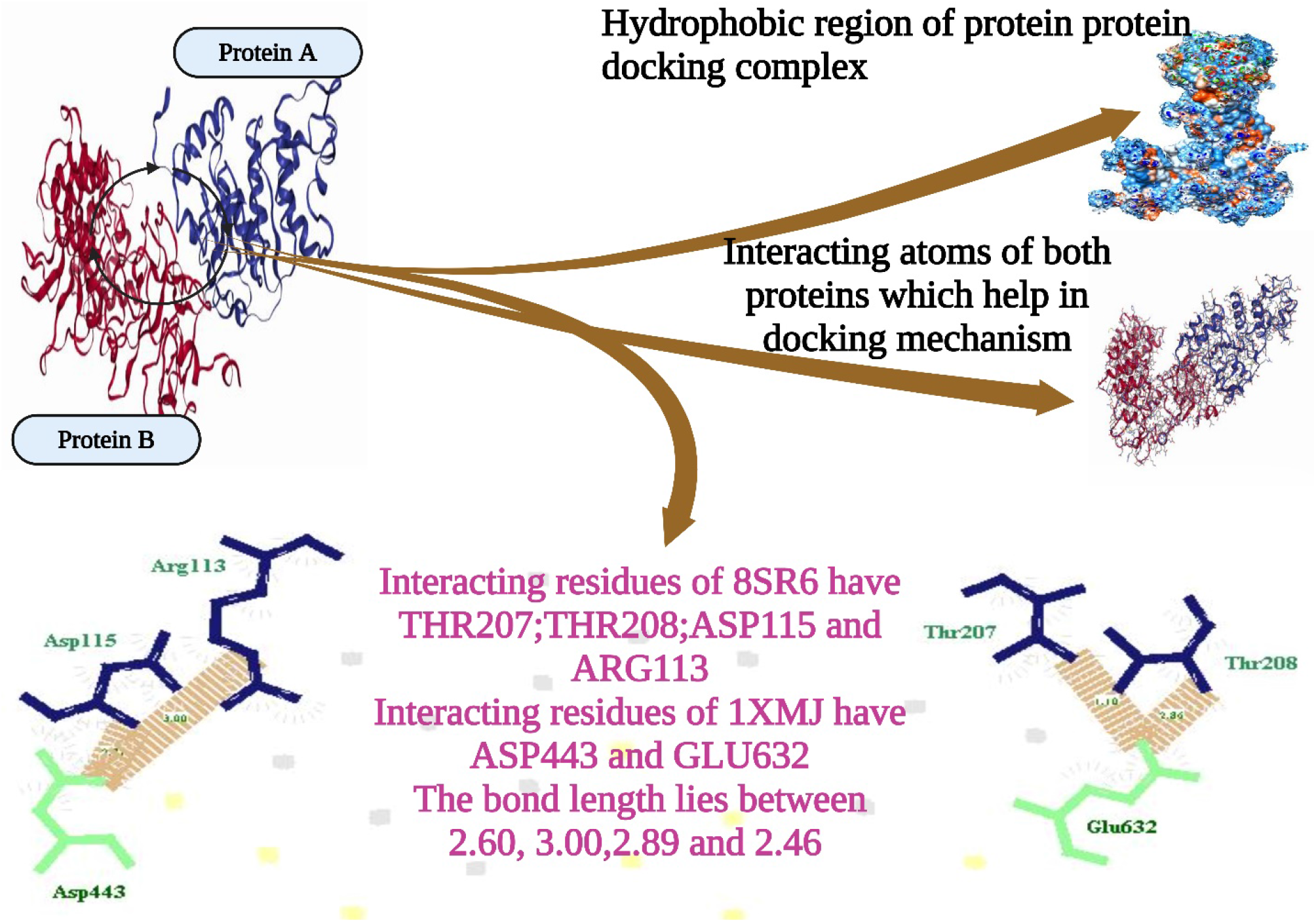
Describe the hydrophobic region, docking phenomenon of both proteins. Both proteins have interact each other with high binding energy and distances

## Conclusion

In this research, in silico siRNA and are designed against the mutated 16S rRNA gene in CRS disease. This research has determined that the joint computational and in vitro methodologies could be valuable for designing the siRNA against transformations of 16SrRNA gene. The in silico methods spare both time and expense, to improve the siRNA possibility to initiate RNAi. In the future, the computational analysis of both siRNA can be further used with the help of in-vitro exploratory techniques to check efficacy and adequacy. Fifteen mRNA were selected in this research for explore targets in CRS disease. All mRNAs have find siRNA regions in their sequence through target finder, No tandem repeats and homology find in all mRNA sequences. All mRNS show siRNA their intra and inter energy and oligomers 5’ to 3’ end sequence region isolated. Check that expression of 16SrRNA gene in CRS disease and explore the regions having presence in whole body. CRS disease is caused by CFTR protein family mutation so we isolate mutant variant of CFTR having accession id is 1XMJ, 16S rRNA protein quotes in 8SR6. Expression of both genes present in whole body network, the Docking interaction of both proteins (1XMJ, 8SR6) shows valuable and predicted results. Proteins protein interaction shows that both protein interact each other and use in future to blockage for CRS disease through computer aided drug designing approach.

## ACKNOWLEDGMENTS

The authors received no specific funding for this work..

